# Integration of locomotion and auditory signals in the mouse inferior colliculus

**DOI:** 10.1101/795872

**Authors:** Yoonsun Yang, Joonyeol Lee, Gunsoo Kim

## Abstract

The inferior colliculus (IC) is the major midbrain auditory integration center, where virtually all ascending auditory inputs converge. Although the IC has been extensively studied for sound processing, little is known about the neural activity of the IC in moving subjects, as frequently happens in natural hearing conditions. Here we show, by recording the IC neural activity in walking mice, the activity of IC neurons is strongly modulated by locomotion in the absence of sound stimulus presentation. Similar modulation was also found in deafened mice, demonstrating that IC neurons receive non-auditory, locomotion-related neural signals. Sound-evoked activity was attenuated during locomotion, and the attenuation increased frequency selectivity across the population, while maintaining preferred frequencies. Our results suggest that during behavior, integrating movement-related and auditory information is an essential aspect of sound processing in the IC.

## Introduction

The inferior colliculus (IC) is the major auditory integration center in the midbrain, where virtually all ascending inputs from the auditory brainstem and the descending cortical inputs converge (Adams, 1979, 1980; Malmierca, 2004; Winer and Schreiner, 2005). The IC plays a critical role in auditory processing, such as representing spectrotemporal features and communication sounds (Egorova et al., 2001; Escabi and Schreiner, 2002; Lesica and Grothe, 2008; Woolley and Portfors, 2013), and sound localization (Bock and Webster, 1974; Schnupp and King, 1997; Lesica et al., 2010; Xiong et al., 2013; Ono and Oliver, 2014). While auditory response properties have been extensively studied, most of neural recordings in the IC have been performed in stationary subjects, and little is known about the IC activity while subjects are engaged in locomotion.

The central nucleus of the IC is considered a predominantly auditory structure. In contrast, the shell region of the IC (the lateral and dorsal cortex) receives non-auditory inputs such as somatosensory inputs (Cooper and Young, 1976; Morest and Oliver, 1984; Coleman and Clerici, 1987; Lesicko et al., 2016) and is thought to perform multi-sensory integration (Aitkin et al., 1978, 1981; Jain and Shore, 2016). The shell region has also been implicated in generating sound-driven behavior by projecting to motor-related regions (Huffman and Henson, 1990; Xiong et al., 2015). Although this multi-modal integration may subserve a range of functions (Gruters and Groh, 2012), modulation of the IC activity during vocalization (Schuller, 1979; Tammer et al., 2004;) or eye movements (Groh et al., 2001; Porter et al., 2006, 2007) suggests that a main role of the non-auditory inputs is providing motor-related information.

Movement adds challenges to auditory processing such as recognizing sounds associated with movement itself or changing spatial relationships with a sound source. In order to accurately detect and localize sounds during movement, it is hypothesized that the auditory system distinguishes self-generated sounds from external ones (Poulet and Hedwig, 2002; Rummell et al., 2016; Schneider et al., 2018) and integrates movement-related signals with auditory information. Recent studies in behaving mice show that movements such as locomotion indeed strongly modulate neural activity in the mouse primary auditory cortex (A1) (Schneider et al., 2014; Zhou et al., 2014; McGinley et al., 2015; Bigelow et al., 2019). Although it has been shown that neurons in A1 receive movement-related signals from sources outside the auditory pathway (Schneider et al., 2014; Nelson et al., 2013; Nelson and Mooney, 2016; Reimer et al., 2016), movement-related modulation is also found in subcortical auditory centers. In the medial geniculate body (MGB) of the thalamus, for instance, sound-evoked activity is attenuated during locomotion (Williamson et al., 2015; Schneider et al., 2014). There is also evidence for motor-related modulation in the auditory brainstem structures during vocalization, licking, and pinna orientation (Suga and Schlegel, 1972; Schuller, 1979; Kanold and Young, 2001; Singla et al., 2017). Evidence of these subcortical modulations suggests that the loci of the integration of movement-related and auditory information are spread out along the auditory pathway, including the IC.

To determine whether and how the neural activity of the IC is modulated during movement, we investigated neural activity of the mouse IC during locomotion. Using an awake head-fixed mouse preparation, we compared the IC neural activity between stationary and walking conditions. Our results demonstrate both spontaneous and sound-evoked activity of IC neurons are strongly modulated during locomotion. Adding to the growing body of evidence of movement-related modulation in the auditory pathway, our results indicate that auditory midbrain neurons receive information about body movement, which may be important for auditory processing during movement and acoustically guided behavior.

## Results

### Spontaneous neural activity of IC neurons is modulated during locomotion

We made extracellular recordings of spiking neural activity of IC neurons in awake head-fixed mice, placed on a passive treadmill. This preparation enabled us to observe the IC neural activity during locomotion. When we compared firing rates between stationary and walking periods, we found that, in the absence of sound stimulus presentation, the firing rates of IC neurons could be strongly modulated during locomotion. Some neurons showed a robust increase in firing during the bouts of locomotion (Figure 1A), while others showed a decrease in firing (Figure 1B). Of 96 recorded IC neurons, 51 neurons (53%) significantly increased their firing during locomotion, while 22 neurons (23%) decreased their firing (Figure 1C). In 23 neurons (24%), firing rates did not significantly differ between the stationary and walking periods. Neurons that increased firing showed positive correlation between the firing rate and the walking speed, whereas neurons that decreased firing showed negative correlation (Figure 1D).

**Figure 1.**
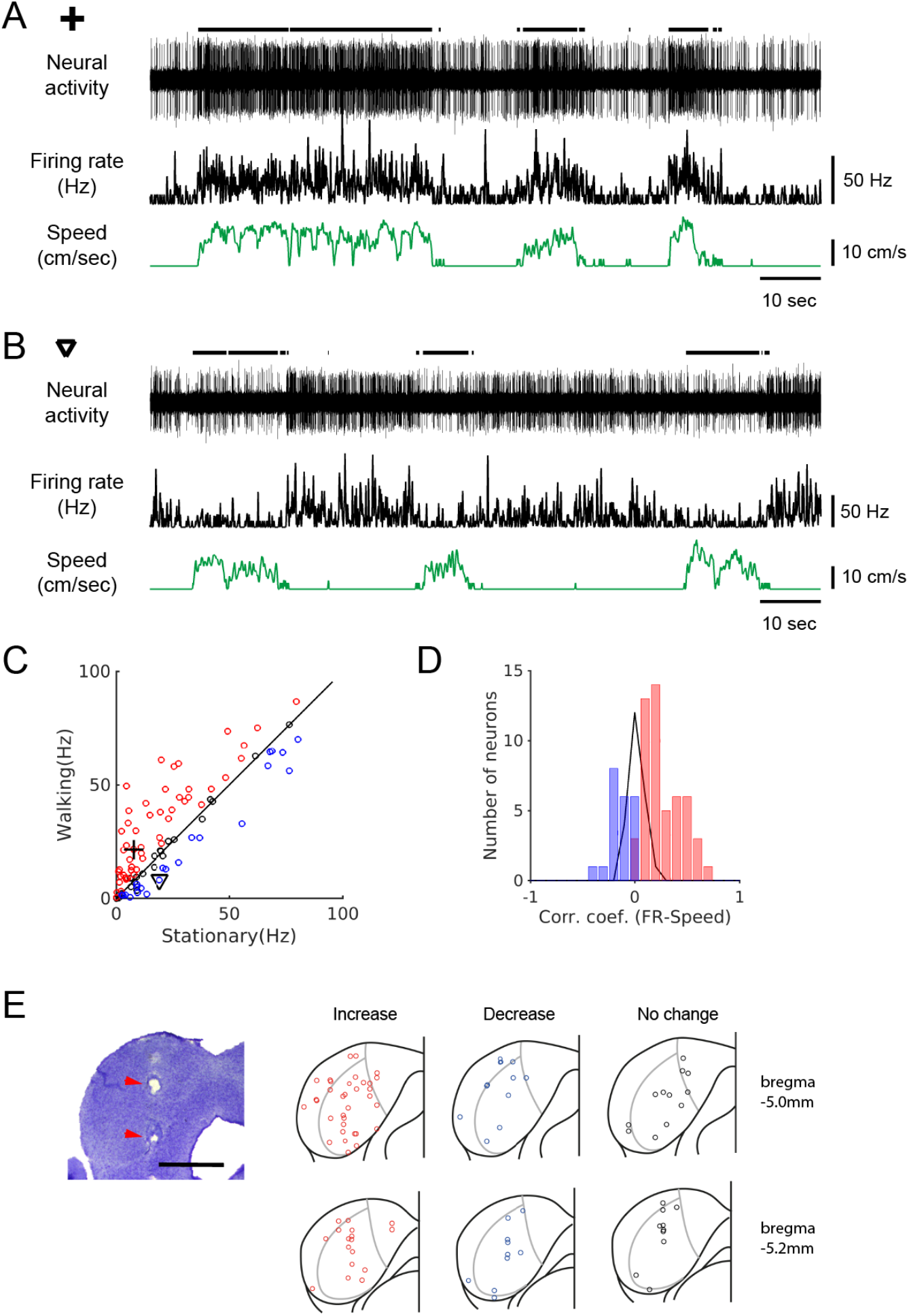
Spontaneous activity of IC neurons is modulated by locomotion. (**A-B**) Example recordings of spontaneous activity in IC neurons. In the neuron in A, the average spontaneous firing rate increased during walking periods (from 7.6 Hz to 21.6 Hz), whereas in the neuron in B, the firing rate decreased during walking periods (from 19.1 Hz to 8.2 Hz). In both cases, the smoothed firing rates (black middle traces) exhibit significant correlations with the speed of the treadmill (green bottom traces) (A: r = 0.59; B: r = −0.24). Thick black lines above the neural records indicate walking periods. (**C**) Population plot comparing average spontaneous firing rates between stationary and walking conditions (n = 96 neurons). Red circles: increase, blue: decrease, black: no significant change in firing. Values for the example neurons in A (cross) and B (triangle) are also shown. (**D**) Histogram of correlation coefficients between smoothed firing rate and speed (color code as in C). (**E**) Photomicrograph of a Nissl section with lesion sites marked with red arrow heads (scale bar = 1 mm), and reconstructed recording locations are shown in 2 transverse sections (5.0 mm and 5.2 mm posterior to the bregma, respectively). Color code as in C.

The subdivisions of the IC – central nucleus, lateral cortex, and dorsal cortex - differ in cytoarchitecture and projection patterns (Morest and Oliver, 1984). For example, the lateral cortex is known to receive somatosensory inputs (Lesicko et al., 2016). To determine whether neurons that showed robust locomotion-related modulation are clustered, we examined the locations of recording sites based on lesions. Anatomical reconstruction of the recording locations did not show any clustering in terms of modulation or the direction of modulation (Figure 1E). Instead, all three types of neurons – increase (red), decrease (blue), or no change (black) - were found across the IC with no clear pattern.

### Neural modulation precedes locomotion onset

When mice walk on a treadmill, the movements generate low intensity sounds. Therefore, the observed locomotion-related modulation in IC neurons could have resulted from auditory responses to the sounds generated by walking. If the firing rate change starts before locomotion onset, however, it would indicate the modulation is not simply due to auditory reafference (Schneider et al., 2014). To determine the timing of modulation with respect to locomotion, we performed locomotion onset-triggered averaging of firing rates across walking bouts of a neuron. In the example neuron shown in Figure 2A, the firing rate begins to increase well before the onset of locomotion. In a different neuron (Figure 2B), a suppression of neural firing occurs before the onset of locomotion. In most of the 30 neurons that yielded significant onset-triggered averages (see Methods), the onset of the firing rate modulation preceded locomotion onset (Figure 2C, n = 30, median latency = −107 msec). These negative latencies were observed regardless of the direction of the modulation (Figure 2D; positive modulation: red bar, n = 22, −118 msec; negative modulation: blue bar, n = 8, −60 msec; Wilcoxon rank sum test, *p* = 0.25). This demonstrates that neural modulation can begin before any detectable movement, which therefore is not simply attributable to auditory responses to walking sounds.

**Figure 2.**
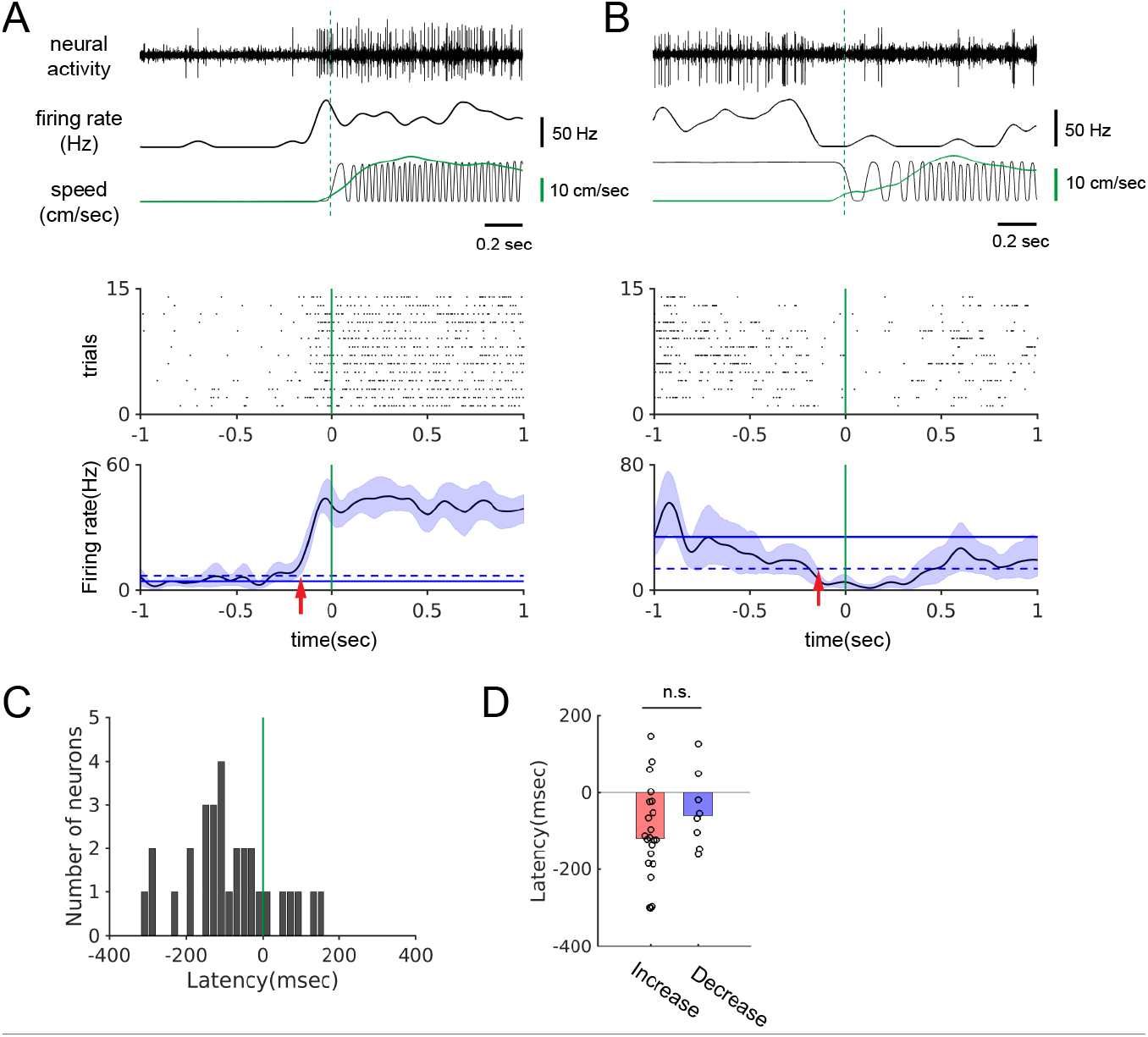
The onset of firing rate modulation precedes locomotion onset. (**A-B**) Top: Example neural records, corresponding smoothed firing rates, and locomotion signals (black: treadmill sensor output, green: speed). Bottom: Spike rasters aligned at the onset of walking and the locomotion onset-triggered averages of firing rates. Blue shades indicate the 95% confidence interval of the average firing rates. Horizontal solid and dashed lines show the average spontaneous firing rate and the two times the standard deviation above (A) or below (B) the average, respectively. Vertical green lines indicate locomotion onset. Red arrows indicate modulation onset. (**C**) Histogram of the neural modulation latencies relative to locomotion onset (n = 30). Time zero indicates locomotion onset. (**D**) Modulation latencies for neurons with increased (red bar, n = 22) and decreased (blue bar, n = 8) firing during locomotion (Wilcoxon rank sum test, *p* = 0.25). Bar graphs show the median latencies.

### Walking sound playback does not mimic neural modulation by locomotion

To directly investigate the contribution of auditory reafference, we examined neural responses to the playback of the recorded walking sounds (n = 25). In our behavioral apparatus, the walking sounds had the sound pressure level of approximately 30 dB or less (see Methods) with sound energy mainly around 4 to 20 kHz (Figure 3A). About ~52% of the recorded neurons did show a significant average firing rate increase during passive playback of the walking sounds (Figure 3B). Across the population, however, the range of the playback-evoked firing rate changes was significantly smaller than that of the changes during locomotion (Figure 3C; *F* test, *p* = 2.7 × 10^−5^).

**Figure 3.**
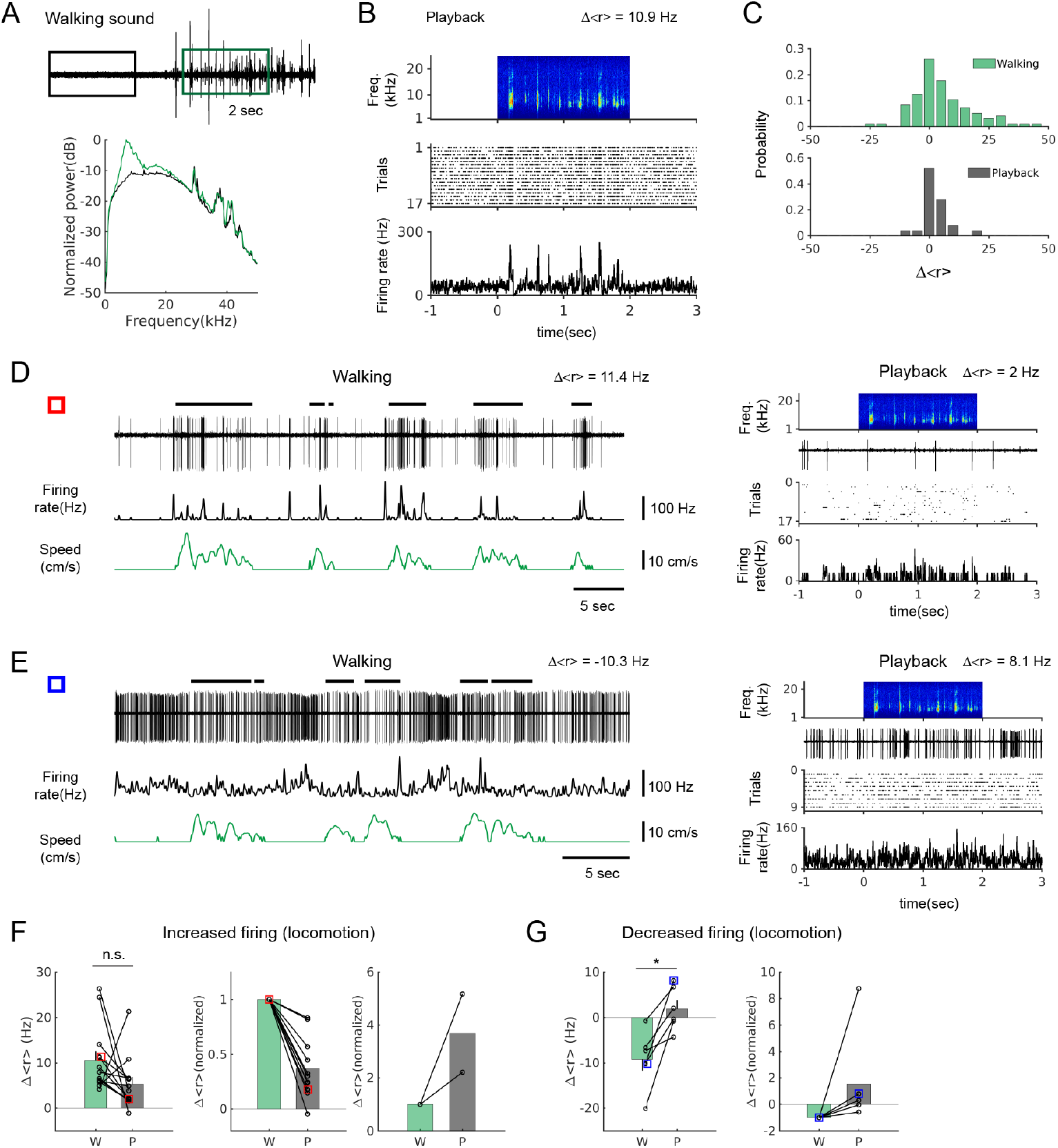
Walking sound playback does not mimic neural modulation by locomotion. (**A**) Top: walking sound recording in sound pressure waveforms (Black and green boxes denote 2 second periods during baseline and walking, respectively, used for power spectrum calculation). Bottom: normalized power spectrum (black trace: baseline, green trace: walking). (**B**) Example neuron that showed a relatively strong response to the playback of walking sounds (Δ<r> = 10.9 Hz). Spectrogram of walking sounds (top), raster plot (middle), and PSTH (bottom) are shown. (**C**) Distributions of average firing rate changes by locomotion (top, n = 96) and by playback (bottom, n = 25). *F* test, *p* = 2.7 × 10^−5^. (**D**) Example neuron that showed a strong positive modulation during locomotion (left; Δ<r> = 11.4 Hz), but a weak response to the playback (right; Δ<r> = 2.0 Hz). (**E**) Example neuron that showed a suppression during locomotion (left; Δ<r> = −10.3 Hz), but an excitatory response to the playback (right, Δ<r> = 8.1 Hz). (**F**) Comparison of average firing rate changes (Δ<r>) during locomotion (green bars) and playback (gray bars) in neurons that increased firing during locomotion (n = 13). Left: Δ<r> comparison (W: walking, P: playback; Wilcoxon signed rank test, *p* = 0.057), Center: Δ<r> normalized to locomotion values (n = 11/13), Right: Δ<r> normalized to locomotion values in neurons with playback response much greater than modulation by locomotion (n = 2/13). Red squares represent the neuron shown in D. (**G**) Same as F, but neurons that decreased firing during locomotion (n = 6). Left: Δ<r> comparison (Wilcoxon signed rank test, *p* = 0.0082), Right: Δ<r> normalized to locomotion values. Blue squares represent the neuron in E.

In 19 of the 25 neurons (76%) that showed significant modulation during locomotion (13 increased; 6 decreased), the firing rate changes during locomotion and playback were compared. As shown in the example neurons (Figure 3D-E), the firing rate change during locomotion was not well mimicked by playback. In neurons that increased firing during locomotion, the overall playback-induced firing rate increase was not significantly different from the increase by locomotion (Figure 3F, left, n = 13; W: 10.6 ± 2.0 Hz; P: 5.3 ± 1.6 Hz, Wilcoxon signed rank test, *p* = 0.057). However, most neurons showed significantly smaller firing rate increase by playback, accounting for ~37% of the modulation by locomotion (n = 11/13, Wilcoxon signed rank test, *p* = 0.00098; Figure 3D and 3F, middle). In the remaining two neurons, the playback-induced firing rate increase was much greater than the increase by locomotion (Figure 3F, right; n = 2/13, 220% and 517%, respectively). Therefore, in this group, the playback response was either much smaller or larger than the modulation by locomotion.

In neurons that decreased firing during locomotion, the firing rate did not similarly decrease during playback (Figure 3E and 3G; n = 6, W: −9.2 ± 2.6 Hz; P: 1.9 ± 1.9 Hz, Wilcoxon signed rank test, *p* = 0.0082). Therefore, in these neurons as well, playback response did not mimic the modulation by locomotion. Taken together, our playback results indicate that although the walking sounds may evoke auditory responses during locomotion, this auditory reafference cannot account for most of the observed firing rate modulation during locomotion.

### Locomotion modulates spontaneous activity in deafened mice

To further substantiate that non-auditory neural signals modulate the IC activity during locomotion, we bilaterally deafened mice by removing the middle ear ossicles and applying an ototoxin (kanamycin) to the cochlea (see Methods; n = 4 mice). The effect of deafening was examined by systematically recording multi-unit responses to broadband noise across the IC. In normal mice, multi-unit responses are evident typically around 30 dB (Figure 4*A*, left and 4B, top). In deafened mice, however, the responses only appeared at 70 dB or higher (Figure 4A, right and 4B, bottom), indicating that the procedure raised hearing thresholds by at least 40 dB. We reasoned that in these mice, it would be highly unlikely that the low intensity walking sounds (< 30 dB, Figure 3) evoke neural responses. As shown in an example neuron from a deafened mouse (Figure 4C), we observed a robust increase in firing during locomotion, while the same neuron did not show any discernible response to broadband sound stimuli (Figure 4D). We observed both increases and decreases in neural firing during locomotion in deafened mice, as was the case in normal hearing mice (n = 34; Figure 4E and 4F). We also anatomically confirmed that our recording sites in deafened mice were in the IC, and as in normal mice, modulated neurons were found across the IC without any spatial patterns (Figure 4G).

**Figure 4.**
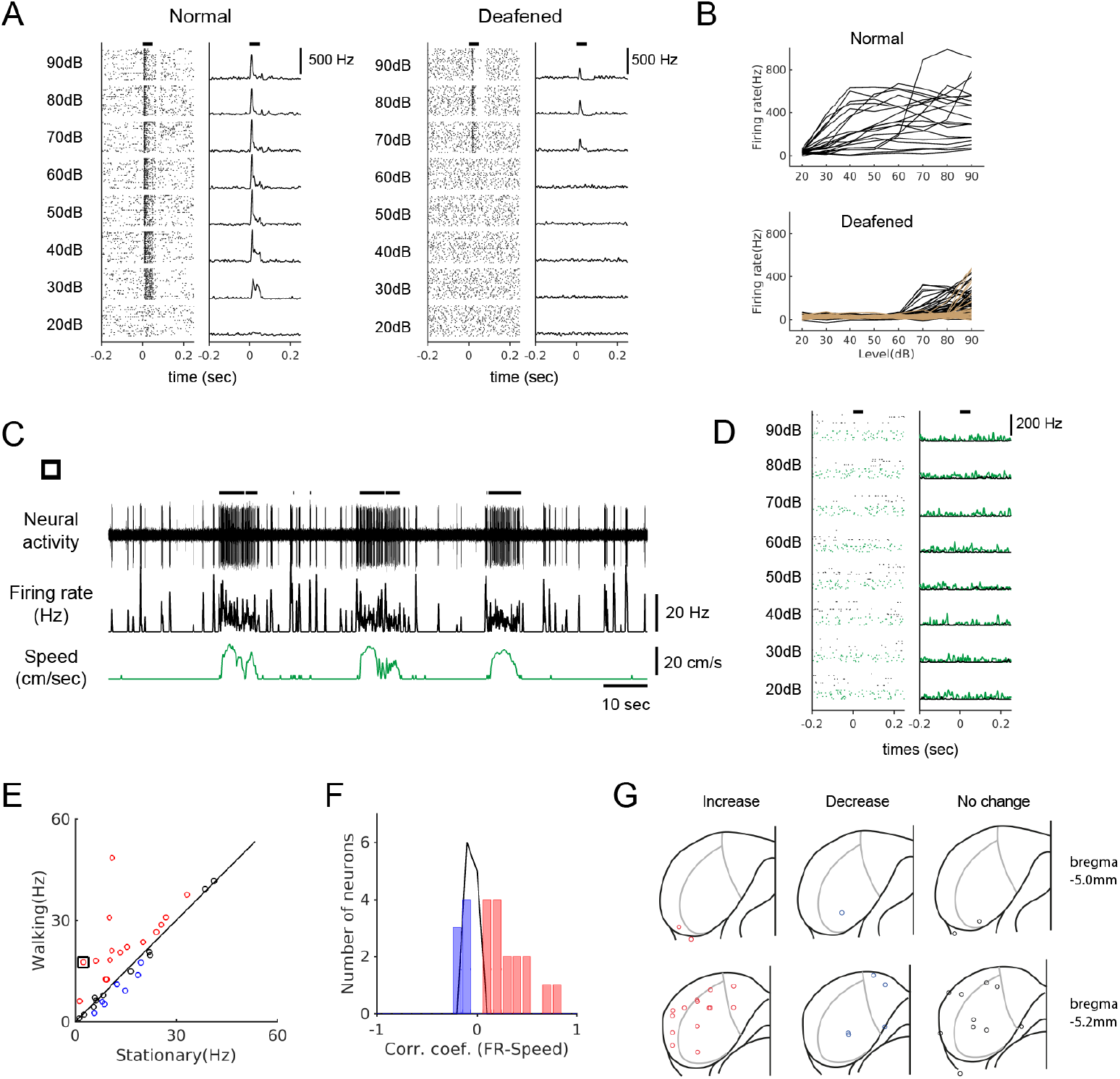
Locomotion modulates spontaneous activity in deafened mice. (**A**) Raster plots (30 trials shown for each level) and PSTHs of multi-unit responses to broadband sound (2-64 kHz, 50 msec duration) in example sites from normal (left) and deafened (right) mice. Horizontal bars at the top denotes the period of stimulus presentation. (**B**) Rate-level functions from multi-unit sites from a normal mouse (top, 20 sites) and two deafened mice (bottom, 40 sites (black) and 57 sites (brown); colors represent different mice, and curves from only two mice are shown for clarity). (**C**) Example neuron from a deafened mouse that showed robust modulation in firing rate during locomotion. Thick black lines above the neural record indicate walking periods. (**D**) The neuron in C did not show any auditory response to broadband noise (black: stationary, green: locomotion). In the raster plots, 30 stationary trials (black dots) and 8-10 walking trials (green dots) are shown for each level. (**E**) Scatter plot comparing the average spontaneous firing rates between stationary and walking conditions in deafened mice (n = 34). Red: increase; blue: decrease; black: no significant change. Black square indicates the neuron in C. (**F**) Histogram of correlation coefficients (color code as in E). (**G**) Reconstructed recording locations as in Figure 1. Color code as in E.

Locomotion-related modulation of spontaneous activity in deafened mice was similar to that in normal hearing mice in several respects. First, the proportions of neurons for each category of firing rate change – increase, decrease, or no change – were comparable (Figure 5A). Second, the degree of modulation, as measured by the modulation index (MI), was comparable between the two groups in both positively (Figure 5B, left; normal: 0.39 ± 0.04, n = 51; deaf: 0.29 ± 0.06, n = 16; *t* test, *p* = 0.18) and negatively modulated neurons (Figure 5B, right; normal: −0.27 ± 0.05, n = 22; deaf: −0.18 ± 0.04, n = 7; *t* test, *p* = 0.28). Third, latencies of modulation onset (Figure 5C; median latencies normal: −107 msec n = 30; deaf: −89 msec, n = 14; Wilcoxon rank sum test, *p* = 0.65) were also comparable. The existence of locomotion-modulated neurons in deafened mice with shared characteristics provides strong evidence that non-auditory signals modulate the activity of IC neurons during locomotion.

**Figure 5.**
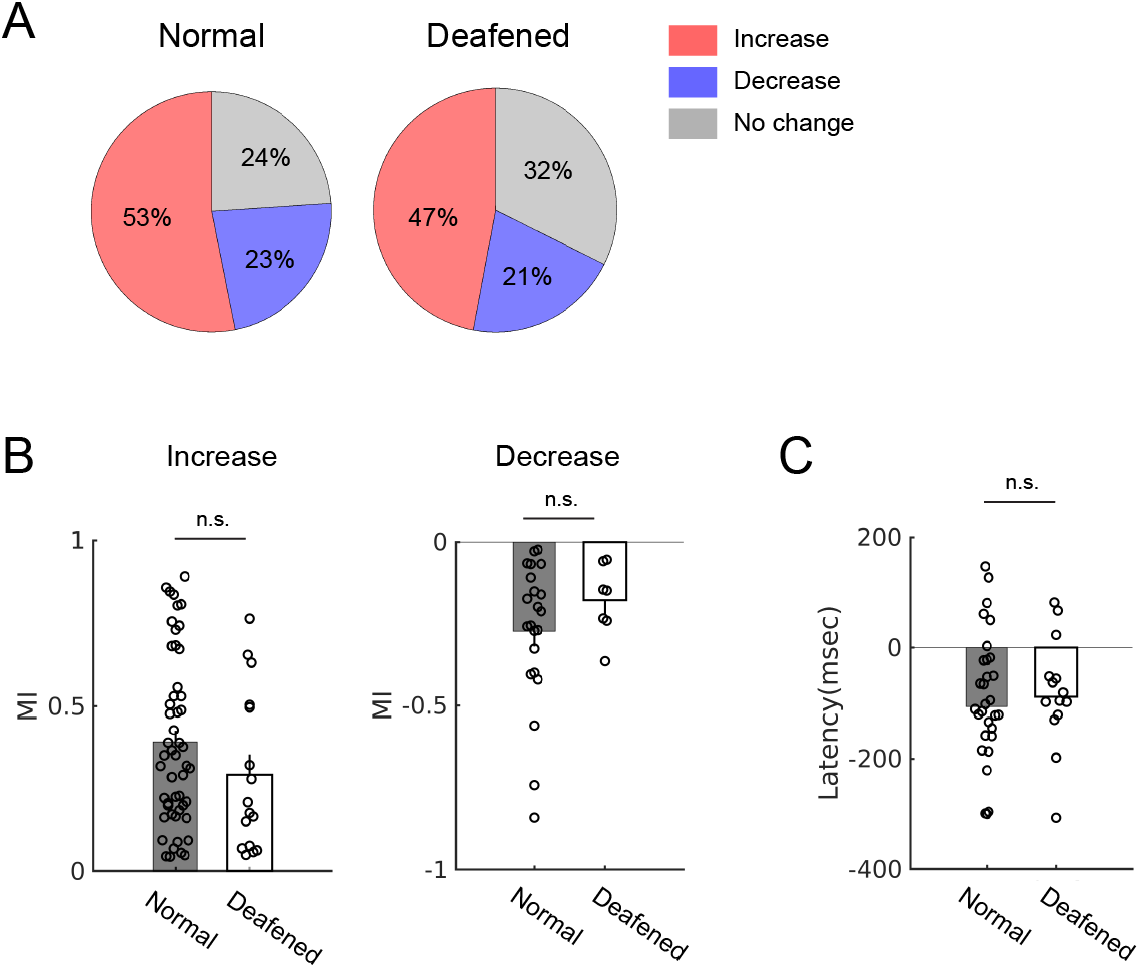
Modulation of spontaneous activity is similar between normal hearing and deafened mice. (**A**) Proportions of the types of modulation: increase, decrease, or no change. Normal: n = 96, deafened: n = 34. (**B**) Modulation index (MI) comparison between normal and deafened mice in neurons that showed positive (left; normal: n = 51; deaf: n = 16; *t* test, *p* = 0.18) or negative modulation (right; normal: n = 22; deaf: n = 7; *t* test, *p* = 0.28). (**C**) Latencies of neural activity modulation onset relative to locomotion onset (normal: n = 30; deafened: n = 14; Wilcoxon rank sum test, *p* = 0.65).

### Sound-evoked activity of IC neurons is attenuated during locomotion

Attenuation of sound-evoked activity during locomotion has been shown in the auditory thalamus and cortex, but sources of subcortical attenuation remain unclear (Schneider et al., 2014; Zhou et al., 2014; Williamson et al., 2015; McGinley et al., 2015). We investigated whether the sound-evoked activity in the IC is also modulated during locomotion by presenting pure tones of different frequencies at 70 dB (Figure 6).

**Figure 6.**
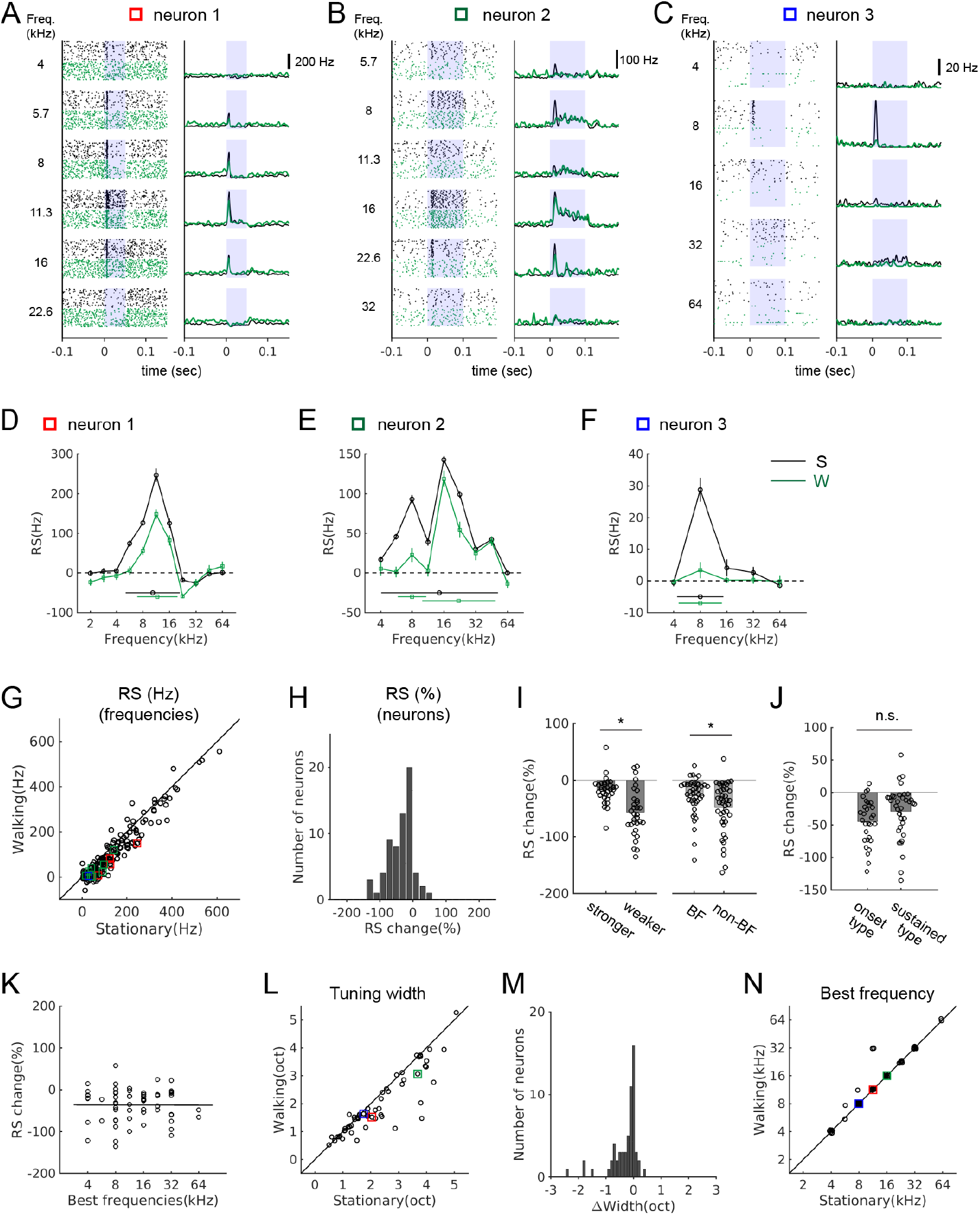
Sound-evoked activity of IC neurons is attenuated during locomotion. (**A-C**) Tone-evoked responses from three example IC neurons. Black spike rasters and PSTHs are from stationary trials, and green from walking trials. In each neuron, the same number of trials is shown for each condition (A: 35 trials; B: 25 trials; C: 92 trials). Blue shades indicate the period of sound stimulus presentation. Squares in different colors indicate the data points corresponding to the examples in the population plots in G, L, and N. (**D-F**) Tuning curves (black: stationary, green: walking) based on response strength (RS, baseline subtracted average firing rates at stimulus onset; see Methods). Tuning widths are shown as horizontal bars in each condition (with the centroid shown as a circle at the center). The tuning curves are from the neurons shown in A-C, respectively. (**G**) Comparison of RS values between the stationary and walking conditions at all frequencies with excitatory responses (178 tones, 65 neurons). Values from the example neurons in A-C are indicated as squares in corresponding colors. (**H**) Histogram of percent changes in RS values across the recorded neurons (n = 65). One sample *t* test for the zero mean, *p* = 1.4 × 10^−10^. (**I**) Comparison of the percent change in RS during locomotion between neurons with weaker and stronger RS (the left two bars; stronger: above the median, n = 33; weaker: below the median, n = 32; *t* test, *p* = 0.0175), or between best and non-best frequencies (BF vs. non-BF) (the right two bars; n = 43 of 65 neurons that had RS values at both best and non-best frequencies; paired *t* test, *p* = 6.98 × 10^−4^). Each data point denotes a neuron. (**J**) Percent change in RS during locomotion in neurons with responses only at the onset vs. neurons with sustained responses (onset type: n = 30; sustained type: n = 35; *t* test, *p* = 0.103). (**K**) Scatter plot of percent change in RS as a function of best frequencies (n = 65; r = −0.0031, *p* = 0.98). (**L**) Frequency tuning widths in octaves for stationary and walking conditions. (**M**) Histogram of tuning width changes (n = 60, one sample Wilcoxon signed rank test, *p* = 3.0 × 10^−7^; mean change = −0.29 octaves). (**N**) Comparison of best frequencies (n = 60). In L-N, neurons that lost all excitatory responses in the walking condition, yielding the tuning width of zero, were not included (n = 5).

We found that most of IC neurons with excitatory tone-evoked responses (measured as response strength, or RS; see Methods) showed significant attenuation during locomotion (72%, n = 47/65; Figure 6A-C and G). Percent changes in evoked response across the population showed a significant shift toward negative values (Figure 6H, n = 65; one sample *t* test, *p* = 1.4 × 10^−10^) with a mean change of −36 ± 5%. Percent attenuation was greater in neurons with relatively weaker responses (below the median) than in neurons with stronger responses (above the median) (Figure 6I, left; −17 ± 4% (stronger, n = 33) vs. −56 ± 7% (weaker, n = 32); *t* test, *p* = 0.0175). Within a neuron, the average attenuation at non-best frequencies were greater than at a neuron’s best frequency (Figure 6I, right; n = 43, −27 ± 5% (BF) vs. −48 ± 7% (non-BF), paired *t* test, *p* = 6.98 × 10^−4^). The greater attenuation of weaker responses within and across neurons may improve signal-to-noise ratio across the population of neurons during locomotion.

To examine whether the degree of attenuation depends on response properties of a neuron, we divided the IC neurons into two response types: one that showed response only at the onset of a stimulus (e.g., Figure 6C; n = 30), and the other that showed sustained response throughout a stimulus (e.g., Figure 6B; n = 35). Although the average attenuation was less in the neurons with sustained response, the difference was not statistically significant (Figure 6J; −45 ± 6% vs. −29 ± 7%; *t* test, *p* = 0.103). The degree of attenuation also did not correlate with best frequencies, indicating the attenuation was global, rather than specific to certain frequencies (Figure 6K, n = 65; r = −0.0031, *p* = 0.98).

Attenuation of evoked activity indicates a decrease in response gain, and this may affect the frequency selectivity of a neuron. To examine this possibility, we constructed frequency tuning curves and quantified tuning widths as the spread around the centroid (see Methods; Escabi et al., 2007; Ono et al., 2017; Figure 6D-F, L, M). In the stationary condition, the tuning widths ranged from 0.5 to 5 octaves with a mean of 2.2 octaves (n = 65). During locomotion, there was a significant decrease in the tuning widths across the population (Figure 6L-M, n = 60, excluding 5 neurons with tuning width of zero in walking condition due to near complete suppression; one sample Wilcoxon signed rank test, *p* = 3.0 × 10^−7^; mean change = −0.29 octaves). In contrast, best frequencies did not change during locomotion in the vast majority of the neurons (n = 60, Figure 6N). Together, these results demonstrate sound-evoked activity in the IC is attenuated during locomotion, and this attenuation significantly sharpens frequency tuning across the population.

### Attenuation of sound-evoked activity can occur independent of spontaneous activity modulation

Having found locomotion-related modulation in both spontaneous and sound-evoked activity, we asked whether there is any relationship between the two, which might indicate shared neural mechanisms. When we compared the modulation of evoked activity across different types of spontaneous activity modulation (increase: n = 39, Figure 7A-B; decrease: n = 8, Figure 7C; no change: n = 18, Figure 7D), evoked activity was attenuated during locomotion in all three groups, and the degrees of attenuation did not differ significantly among the groups (Figure 7E; increase (red): −31 ± 6%, n = 39; decrease (blue): −46 ± 15%, n = 8; no change (gray): −45 ± 10%, n = 18; one way ANOVA, *p* = 0.32). Thus, attenuation of evoked activity during locomotion can occur independent of the modulation of spontaneous activity.

**Figure 7.**
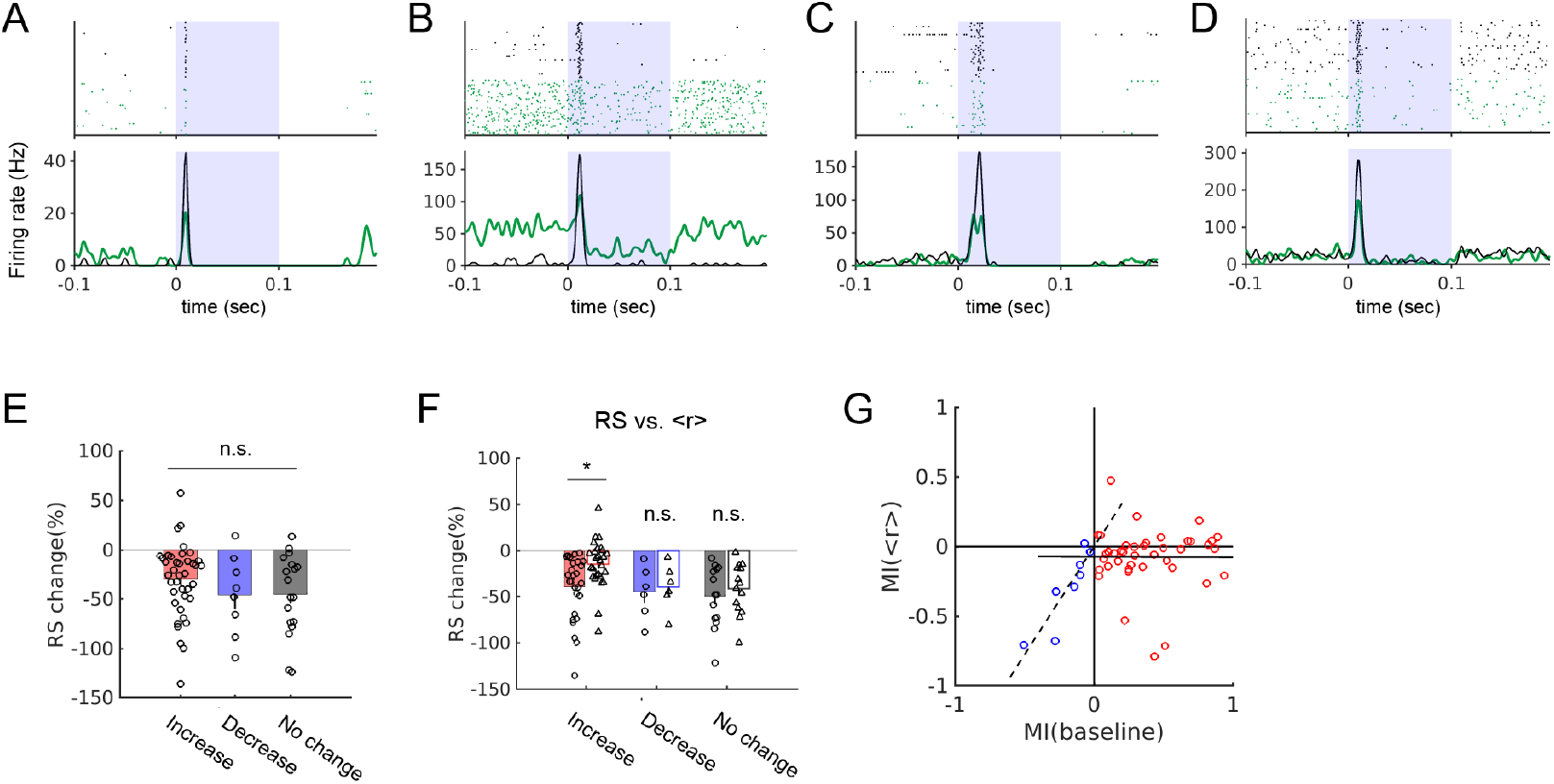
Relationships between the locomotion-related modulation of spontaneous and tone-evoked activity. (**A-D**) Tone-evoked responses in four example neurons with different types of spontaneous activity modulation during locomotion: increase (A-B), decrease (C), or no change (D). In raster plots and PSTHs, black indicates stationary and green indicates walking trials. (**E**) Attenuation of evoked activity (RS) in the groups with different types of spontaneous activity modulation. Red: increase, n = 39; blue: decrease, n = 8; gray: no change, n = 18. (**F**) Attenuation of evoked rates with or without baseline rate subtraction (RS vs. <r>) in the three groups in E. These comparisons were made only in neurons that showed significant evoked response attenuation (*t* tests, red: n = 27 of 39, *p* = 0.0035; blue: n = 6 of 8, *p* = 0.72; gray: n = 14 of 18, *p* = 0.47). Color coding is the same as in E. (G) Pairwise scatter plot of modulation index (MI) for spontaneous rates and <r>. MI(spontaneous) and MI(<r>) values are shown for neurons with positive spontaneous activity modulation (red; n = 39; r = −0.0044, *p* = 0.98) and negative spontaneous activity modulation (blue; n = 8; r = 0.899, *p* = 0.0024). Corresponding linear fits are shown in solid (for red circles) and dashed (blue circles) lines.

In the group with increased spontaneous activity, we also examined whether the attenuation in the evoked activity is simply due to the increased baseline firing rate. In neurons that showed attenuation (n = 27/39), the percent change in firing rates during sound presentation without baseline subtraction (denoted as <r>) was significantly smaller than the change in RS. Therefore, in this group, the modulation of spontaneous activity played a significant role in our measurement, but it didn’t entirely account for the attenuation (Figure 7F, red bars, −40 ± 7% vs. −15 ± 5%, *t* test, *p* = 0.0035, n = 27 of 39). In the group with decreased spontaneous activity, the change in RS did not differ from the change in <r> (Figure 7F, blue bars, −45 ± 12% vs. −40 ± 10%, *p* = 0.72, n = 6 of 8). This was also true, as expected, for the group with no significant modulation of spontaneous activity (Figure 7F, gray bars, −50 ± 9% vs. −42 ± 7%, *p* = 0.47, n = 14 of 18).

Finally, we asked whether there is a relationship between the magnitude of spontaneous activity modulation and evoked activity modulation. For this analysis we used modulation index (MI) for the spontaneous rates and <r>. In the group with increased spontaneous activity, the magnitude of the increase did not correlate with the modulation in <r> (n = 39, solid line, r = −0.0044, *p* = 0.98; Figure 7G). In contrast, in the group with decreased spontaneous activity, the magnitude of decrease showed a significant positive correlation with the modulation in <r> (n = 8, dashed line, r = 0.899, *p* = 0.0024; Figure 7G). These results suggest that the suppression of spontaneous and evoked activity may share common neural mechanisms, whereas the increase in spontaneous activity during locomotion may result from distinct sources.

## Discussion

By recording single unit activity in the IC of behaving mice, we found that locomotion can bidirectionally modulate spontaneous activity of IC neurons. The modulation preceded locomotion onset, was not mimicked by playback of sounds generated by locomotion, and occurred also in deafened mice. Furthermore, sound-evoked activity was attenuated during locomotion, and frequency selectivity increased. Our results reveal locomotion-related neural signals at this relatively early stage of the auditory pathway. The prevalence of the clear movement-related signals suggests that multi-modal integration of movement and sound is an essential part of sound processing in the IC.

### Modulation of neural activity by locomotion in the IC

We found that in ~50% of the recorded IC neurons, spontaneous firing rates increased during locomotion, while in ~25%, firing rates decreased. This modulation of spontaneous activity contrasts with the findings of prior studies in the MGB, where spontaneous firing did not significantly change during locomotion (Zhou et al., 2014; Williamson et al., 2015; McGinley et al., 2015). Our results show that the modulation during locomotion can be strong and is wide spread in the IC, so it is unlikely that the altered activity does not propagate to the MGB. One plausible explanation of the discrepancy is that the modulation of IC spiking activity primarily evokes subthreshold responses in thalamic neurons, making it difficult to detect in terms of firing rate changes. However, subthrehold inputs could still modulate sound-evoked activity (Jain and Shore, 2006) and could thus contribute to the suppression of evoked activity shown in the MGB during locomotion (Williamson et al., 2015; McGinley et al., 2015).

To better understand the impact of the modulation on the IC circuits, it will be important to determine whether directions or degrees of modulation differ in different cell types. For example, in A1, a major mechanisms of locomotion-related modulation is feedforward inhibition in which the motor cortex inputs excite fast-spiking inhibitory interneurons, which in turn suppresses excitatory neurons (Schneider et al., 2014). In contrast to cortical neurons (Niell and Stryker, 2010), in the IC, recordings from genetically labeled neurons have shown that excitatory and inhibitory neurons cannot be distinguished based on spike waveforms (Ono et al., 2017). Furthermore, defining cell types beyond excitatory and inhibitory neurons remains a challenge in the IC, and different cell types and their connectivity are only beginning to be understood (Oliver et al., 1994; Ito et al., 2009; Beebe et al., 2016; Chen et al., 2018; Goyer et al., 2019; Schofield and Beebe, 2019). Making recordings from defined IC cell types in the future would enable investigations of questions such as whether excited neurons vs. suppressed neurons are different types of neurons in the IC, or how the network as a whole is modulated.

While spontaneous activity was modulated bidirectionally, evoked activity was attenuated in the vast majority of IC neurons. This attenuation was general rather than selective in that it occurred across neurons with different best frequencies, both at best and non-best frequencies of a neuron, and regardless of temporal response patterns. In contrast to spontaneous activity, the attenuation of evoked activity is consistent with the findings in the MGB. Although it is not straightforward to compare the degree of modulation, comparable amount of attenuation of evoked activity has been shown in the MGB (~15-25% in Williamson et al., 2015; ~50% in McGinley et al., 2015). Therefore, a significant portion of the attenuation observed in the MGB already occurs in the IC.

Interestingly, the attenuation of evoked activity led to a decrease in frequency tuning widths, indicating increased neural selectivity for sound frequency (Fig. 6*L-M*). Across the population, we observed an average tuning width decrease of ~0.3 octaves. Behavioral studies indicate that mice can discriminate at least ~10% difference (~0.14 to 0.15 octaves) in frequency (Ehret, 1975; Clause et al., 2011; de Hoz and Nelken, 2014; Guo et al., 2017). Thus, the observed changes in tuning widths is large enough to have significant impact on the sound frequency processing. The attenuation of sound-evoked activity may decrease the sensitivity to sounds, as supported by poorer performance on tone detection tasks during locomotion (McGinley et al., 2015; Schneider et al. 2018). On the other hand, the observed increase in frequency selectivity in IC neurons may improve frequency discrimination (Aizenberg et al., 2015; Carcea et al., 2017; Guo et al., 2017).

### Potential sources of modulation during locomotion

It is likely that the observed modulation has multiple neural sources, as indicated by the result that evoked modulation can occur independent of spontaneous modulation (Figure 7). One obvious candidate, however, is somatosensory feedback during movement. The IC receives inputs from the somatosensory areas in the brainstem and the cortex (Cooper and Young, 1976; Aitkin et al., 1978; Robards, 1979; Coleman and Clerici, 1987; Kunzle, 1998; Zhou and Shore, 2006; Lesciko et al., 2016; Olthof et al., 2019), and sound-evoked activity is modulated by concurrent stimulation of somatosensory afferents (Aitkin et al., 1978; Jain and Shore, 2006). While the exact functions of the somatosensory inputs during behavior still remain poorly understood, our results suggest that a major function of the somatosensory inputs to the IC is to provide feedback about ongoing movement.

The somatosensory projections predominantly innervate the lateral cortex of the IC (Kunzle, 1998; Zhou and Shore, 2006; Lesicko et al., 2016), but our anatomical reconstruction of recording locations indicate modulated neurons are distributed throughout the IC (Figure 1E). Neurons in the central nucleus may receive somatosensory inputs from the lateral cortex via connections within the IC (Rockel and Jones, 1973; Coleman and Clerici, 1987; Jen et al., 2001; Chen et al., 2018), or from the dorsal cochlear nucleus (Li and Mizuno, 1997; Oertel and Young, 2004; Shore 2005; Goyer et al., 2019), which receives somatosensory feedback during behavior (Kanold and Young, 2001; Singla et al., 2017). Descending inputs from the somatosensory cortex, spread out across the IC subdivisions, may also play a role (Olthof et al., 2019). Regardless of the precise neural mechanisms, our results suggest that movement-related somatosensory feedback signals are more wide spread across the IC than previously hypothesized.

Our timing analysis shows a substantial fraction of IC neurons change their firing prior to (~100 msec) movement onset (Figure 2). This relative timing is compatible with efference copy of motor signals, or neuromodulatory signals associated with movements, which has been described in A1 (Nelson et al., 2013; Schneider et al., 2014; Nelson and Mooney, 2016; Reimer et al., 2016). In fact, the IC receives inputs from a number of motor-related regions and neuromodulatory centers likely associated with locomotion. First, IC neurons receive inputs from the superior colliculus, a highly multi-modal area, whose neurons are also modulated by locomotion (Ito et al., 2017), and the motor cortex (Olthof et al., 2019). Second, the IC receives inputs from midbrain cholinergic neurons in the peduncular pontine nucleus (Farley et al., 1983; Motts and Schofield, 2009), which is part of the midbrain locomotion region (Lee et al., 2014; Caggiano et al., 2018). Third, the IC receives noradrenergic inputs from the locus coeruleus (Klepper and Herbert, 1991; Hormigo et al., 2012), which are likely to be active during locomotion (Reimer et al., 2016). Therefore, IC neurons could receive rich information about body movement via both efference copy like signals from motor-related regions and somatosensory feedback. These signals may work together to inform IC neurons of ongoing movements to be integrated with sound processing.

### Implications for sound processing and sound-guided behavior

In the visual system, visual responses are stronger during locomotion, and this increased gain enhances visual function (Niell and Stryker, 2010; Mineault et al., 2016; Dadarlat and Stryker, 2017; Ito et al., 2017). In contrast, in the auditory pathway, it has been consistently observed that auditory responses are suppressed during motor behavior, including locomotion (Creutzfeld et al., 1989; Eliades and Wang, 2013; Singla et al., 2017; Schneider and Mooney, 2018). This suppression is thought to help maintain sensitivity to sounds by preventing desensitization and help distinguish self-generated and external sounds (Poulet and Hedwig, 2002; Schneider et al., 2018). The general attenuation of auditory response we observed in the IC is in line with this sound processing strategy. However, the integration of auditory and movement-related signals in the IC may have additional functional consequences such as a trade-off between detection and discrimination, as suggested by the ways IC tuning curves changed.

The movement-related signals in the IC may also enable rapid control of behavioral response to a sound source. The IC has been implicated in mediating acousticomotor behavior by conveying auditory information to motor-related areas, such as the superior colliculus and the periaqueductal gray (Huffman and Henson, 1990; Xiong et al., 2015). Integrating neural signals related to body movement and posture at the level of auditory midbrain could help localize a sound source and generate rapid response toward or away from it. Through this multi-modal integration, the IC may take part in the midbrain sensory-to-motor circuits that allow rapid control of sound-driven behaviors.

## Materials and Methods

### Animals

All experiments performed were approved by the Institutional Animal Care and Use Committee of Sungkyunkwan University in accordance with the National Institutes of Health guidelines. Neural recordings have been performed in C57BL/6 mice (n = 15 mice) or a transgenic mouse line (VGAT-ChR2-EYFP line with C57BL/6J background; n = 10 mice) of both sexes, aged 6-10 weeks. In the current study, we did not make use of the transgene expression in the transgenic line. We did not find differences in the degree of modulation in spontaneous or evoked activity between the two strains.

### Headpost surgery

For neural recordings in a head-fixed preparation, a custom metal headpost was cemented to the skull. Mice were positioned in a stereotaxic device (Kopf Instrument) and anesthetized using Isoflurane (1-4%) delivered via a vaporizer (DRE Veterinary). An eye ointment was applied to keep the eyes from drying. A small amount of lidocaine (2%) was injected under the skin overlying the skull, and an incision was made to expose the skull. The connective tissues were gently removed, and the skull surface was allowed to dry. The head was positioned such that difference in the dorso-ventral coordinates of the bregma and lambda was less than 100 μm. A small ground screw was cemented toward the rostral end of the exposed skull surface. For future craniotomy over the inferior colliculus (IC), markings were made around 5 mm posterior to the bregma. A metal headpost was positioned not to obstruct future access to the IC and was secured using dental cement (Super-bond C&B).

### Neurophysiology

Mice were acclimated to head-fixing and walking on a passive disc-type treadmill for 2-4 days (one 30 min session per day) prior to neurophysiological recordings. On the day of neural recording, a cranial window (~1 mm) was made over the IC under isofluorane anesthesia (1-4%). The window was covered with Kwik-Cast (WPI) and mice were allowed to recover for at least 2 hours. Recordings were made from the IC (AP ~5.0 mm posterior to the bregma, ML ~0.4-1.8mm from the midline) using either a single tungsten electrod or a linear array of tungsten electrodes (~5 MΩ, FHC). The electrodes were controlled by a single-axis motorized micro-manipulator (IVM Mini, Scientifica). Neural signals were acquired using a 16-channel headstage (RHD2132, Intan Technologies) and Open Ephys data acquisition hardware and software. Spiking activity was band-pass filtered (600-6000 Hz) and digitized at 30 kHz. Locomotion on a treadmill was detected using a rotary encoder (Scitech Korea), and the output voltage signal was recorded as analog and digital inputs for further analysis.

### Sound stimuli

Pure tone stimuli with a sampling rate of 400 kHz were generated in MATLAB (MathWorks), and presented at 70 dB SPL using a D/A converter (PCIe-6343, National Instruments), a power amplifier (#70103, Avisoft), and an ultrasonic speaker (Vifa, Avisoft). The sound system was periodically calibrated for each tone stimulus frequency using a ¼” microphone (Bruel & Kjaer 4939). During neurophysiological recordings, the speaker was placed 15 cm from the animal’s right ear at 45 degrees from the body midline. In initial recordings, the tone stimulus set consisted of frequencies between 4 kHz and 64 kHz in one octave steps (duration: 100 msec; on and off ramps: 1 msec; presented pseudorandomly at 2 Hz). In subsequent experiments, to better estimate frequency tuning, tone stimuli were presented at frequencies between 2 kHz and 64 kHz in half octave steps (duration: 50 msec; on and off ramps: 1 msec; presented pseudorandomly at 4 Hz). Each tone stimulus was repeated at least 20 times, but typically much more (mean: 131 trials; range: 20-320). Broadband noise stimuli (2-64 kHz) for assessing hearing thresholds were 50 msec long (5 msec on and off ramps) and presented at 20 to 90 dB SPL in 10 dB steps in pseudorandom order. Each sound level was repeated at least 50 times.

Sounds generated by locomotion on our treadmill were recorded by placing a microphone (CM16/CMPA, Avisoft) close to the legs. The recorded sound signal was bandpass filtered (1kHz-25kHz), and subjected to a noise reduction procedure to reduce the baseline noise using Audition (Adobe). The sound level of walking sounds was estimated by comparing recorded walking sounds with a series of recorded playbacks at different levels (20 to 50 dB in 5 dB steps). Recorded playback at 30 dB had an RMS value similar but slightly higher than that of the recorded walking sounds. Therefore, a representative 2 second recording was used as a playback stimulus. The stimulus was presented at 30 dB SPL and repeated at least 20 times.

Sound stimuli were presented using a custom-written stimulus presenter program written in Python 2.7 (by Jeff Knowles; https://bitbucket.org/spikeCoder/kranky), which communicated with Open Ephys GUI software (http://www.open-ephys.org/gui).

### Deafening

To prevent mice from hearing sounds generated by locomotion, a bilateral deafening procedure was performed. Mice were anesthetized by intraperitoneal injection of Ketamine/ Xylazine (100mg/10mg per kg). A small incision was made ventral and posterior to the pinna. To expose the auditory bulla that surrounds the middle ear cavity, the overlying tissue was gently spread using fine forceps. An opening was made in the bulla to visualize the ossicles and the cochlea. The ossicles were removed and kanamycin drops (1 mg/ml) were applied 3-4 times at the oval window of the cochlea in an attempt to induce further hair cell damage. The middle ear cavity was filled with gelfoam, the overlying tissue was closed, and the skin was sutured. Mice were given analgesics (meloxicam, 5mg/kg) and allowed to recover for 2-3 days before receiving headpost surgery for neural recording.

### Data analysis

Spikes were detected and sorted offline using commercial spike sorting software (Offline Sorter v4, Plexon). Detected spike waveforms were clustered using PCA, and clusters with a clear separation in PC space was taken as single units. A refractory period of 0.7 msec was imposed, and the rate of refractory period violation was required to be less than 0.5%.

Spontaneous activity was measured either from the baseline period that preceded a tone presentation (0.1 or 0.2 sec) or from 1-sec segments of a continuous recording of spontaneous activity (> 200 sec). Periods of locomotion was determined by thresholding the speed of the treadmill at 2 cm/sec, obtained by smoothing the digital recordings of the treadmill sensor output (200 msec hanning filter; a transition from low to high, or vice versa, corresponded to 0.26 cm). During the stationary periods, occasional short blips of movement occurred, but otherwise, the speed was zero. The segments of spontaneous activity was assigned to either stationary or walking condition, and only the segments that occurred entirely during one of the behavioral conditions were included for analysis.

To analyze the timing of neural modulation relative to movement onset (Figure 2), walking periods with clear onsets were identified. Then, for each walking period, the onset was defined as the time when the treadmill sensor output changed by 2% (corresponding to < 1 mm of travel) of the maximum range (2.5V). Once locomotion onsets (= onset(L)) were defined, onset-triggered averaging of the smoothed firing rate was performed. The onset of neural activity modulation (= onset(M)) was defined as the time when the 95% confidence interval of the onset(L)-triggered firing rate first deviated from the average stationary rate by 2 times the standard deviation. The confidence intervals were obtained using bootstrap resampling of the onset(L)-triggered firing rate segments. Latency of modulation was defined as the time of onset(M) - the time of onset(L). Clear onset(L)-triggered averages and modulation onset latency could be defined in a subset of neurons with significant modulation (normal: n = 30 of 63 modulated neurons; deaf: n = 14 of 23 modulated neurons) due to not enough locomotion onsets or relatively weak modulation.

To assess the effect of deafening, multi-unit responses to broadband noise (2-64 kHz) were examined across the IC for a range of sound levels in 10 dB steps (20-90 dB). Multi-unit spike times were obtained by thresholding the neural recordings at 3 times the standard deviation of the baseline noise. Rate-level curves were constructed based on the peak evoked firing rates (Figure 4A-B). Rate-level curves were obtained from a few tens of multiunit sites for each deafened mouse (4 mice, 43 ± 13 sites per mouse).

To compare the degrees of modulation between the normal hearing and deafened mice (Figure 5B), a modulation index was used: MI = [<r>(walk)–<r>(stationary)] / [<r>(walk) +<r>(stationary)], where <r> represents the average firing rate and MI values varied between −1 and +1 (Rummel et al., 2016). Because deafening decreased spontaneous firing rates (21.9 ± 2.2 Hz vs. 14.3 ± 1.8 Hz), MI allowed us to compare the relative changes by locomotion. MI was also used to test whether modulations of spontaneous activity and evoked activity were correlated (Figure 7G).

Tone-evoked activity was analyzed in neurons with excitatory response to at least one of the presented tone frequencies. A significant response to a tone stimulus was determined using a paired *t* test between the firing rate during the baseline period preceding the tone and the firing rate during a response window. A response window was defined around the time of the peak of the smoothed peristimulus time histogram (PSTH) (from 5 msec before and to 7 msec after the peak; smoothing by a Gaussian function with the standard deviation of 2 msec) at a neuron’s best frequency (the tone frequency with the greatest peak response). The same response window was used for all stimuli in a given neuron. Tone-evoked activity was quantified as the average firing rate during a response window minus the spontaneous firing rate (Response Strength or RS; Doupe, 1997). Spontaneous firing rate was measured during a 100 or 200 msec period preceding each stimulus presentation. Evoked trials were assigned to either stationary or walking condition as in the spontaneous activity described above. Only neurons that had at least 5 repeats for a tone stimulus in both conditions were included (Stationary: 131 ± 47 trials per stimulus; Walking: 40 ± 26 trials per stimulus). Modulation of evoked activity was quantified as percent change in RS (100*[RS(walking) - RS(stationary)]/[RS(stationary)]). Percent change in RS for a neuron was defined as percent change in RS summed over all responsive tone frequencies. Modulation analysis using a fixed 15 msec response window starting 5 msec after the sound onset yielded similar results.

Tuning curves were constructed from the average evoked firing rates (RS) at different tone frequencies. To quantify tuning widths, tuning curves were first linearly interpolated between neighboring frequencies (100 points), and tuning widths were expressed as four times the second moment about the centroid, measured in octaves (Escabi et al., 2007; Ono et al., 2017). In rare cases (n = 2) where multiple tuning width segments occurred due to a non-responsive tone in the middle, the widths were added up minus the overlap.

### Histology

At the end of a recording session, small lesions were made by applying current (30 μA, 10 sec) through recording electrodes. Animals were transcardially perfused using PBS followed by 4% paraformaldehyde. Brains were post-fixed for at least a day and were cryoprotected in 30% sucrose before they were cut on a cryostat. Sections were cut at 40-μm thickness, mounted on slides, and processed for Nissl staining. Recording locations were estimated based on the locations of the lesion.

### Statistical analysis

All statistical analysis was performed in MATLAB. The significance level was α = 0.05 except for spontaneous activity modulation, where α = 0.01. Normality was assessed using Lilliefors test (lillietest function in MATLAB), and when data significantly deviated from normal distribution, non-parametric tests were used. Results are presented with mean ± SEM unless otherwise noted.

Statistical significance of the modulation in spontaneous firing was determined using the mean firing rates from recording segments (see *Data analysis* above) during which no sound stimuli were presented. The spontaneous firing rates of the analyzed segments were generally not normally distributed, so the significance of the modulation was determined using a permutation test. In each permutation, segments of spontaneous activity from the stationary or walking groups were combined and then randomly assigned to two groups with the original sample sizes. A distribution of the differences of the means between the two groups was obtained from 1000 permutations. A two-tailed *p* value was obtained by calculating the probability that permuted differences were more extreme than the sample mean difference. For each neuron, *p* < 0.01 was considered a significant modulation. Based on this analysis, neurons were categorized into 3 groups: neurons with increased or decreased spontaneous activity, or with no significant change (normal: n = 96 neurons from 21 mice; deafened: n = 34 neurons from 4 mice; Figures 1C-E, 3F-G, 4E-G, 5A-B, and 7E-G).

In Figure 2D, a Wilcoxon rank sum test was performed to determine if median latencies of neural modulation relative to locomotion onset differed significantly between the neurons with increased (n = 22) and decreased (n = 8) spontaneous firing rates during locomotion. Similarly in Figure 5C, latencies were compared between normal (n = 30) and deafened mice (n = 14).

In Figure 3, whether a neuron showed significant response to walking sound playback was determined using a paired *t* test between the baseline firing rate and the firing rate during the playback. For this analysis, only playback responses from stationary trials were used (17 ± 5 trials). In Figure 3C, to determine whether the variances of the mean firing rate changes (Δ<r>) during locomotion and playback differed significantly across the population, a two sample *F* test was performed (walking: n = 96; playback: n = 25). Randomly selecting 25 neurons from the walking group did not change the result of the statistical test. In Figure 3F and 3G, Wilcoxon signed rank tests were performed for paired comparisons of the mean firing rate changes (Δ<r>) during locomotion and playback in 19 modulated neurons out of 25 with both measurements (Figure 3F: n = 13 neurons in which locomotion increased firing; Figure 3G: n = 6 neurons in which locomotion decreased firing).

In Figure 5B, to determine whether the degrees of modulation differed significantly between the normal and deafened mice, *t* tests were performed on MI values in the neurons with increased (Figure 5B, left; normal: n = 51 of 96 neurons; deafened: n = 16 of 34 neurons) or decreased spontaneous firing (Figure 5B, right; normal: n = 22 of 96 neurons; deafened: n = 7 of 34 neurons). Statistical comparisons using percent changes in firing rate yielded similar results.

In the analysis of the modulation of evoked activity (Figure 6), of the 67 neurons in which tone stimuli evoked excitatory responses, 65 were included in the analysis. Two of the 67 neurons that had unreliable percent change values due to RS values close to zero were excluded from the analysis. In Figure 6H, one sample *t* test was performed to determine whether the population mean of percent changes in RS was significantly different from zero (n = 65). In Figure 6I (two bars on the left), the 65 neurons were divided into two groups: those with relatively stronger (above the median, n = 33) and weaker responses (below the median, n = 32). Then a two sample *t* test was performed to determine whether RS change differed between the two groups. In Figure 6I (two bars on the right), percent changes in RS were compared between the best and non-best frequencies of a neuron using a paired *t* test (n = 43 of 65 neurons with multiple frequencies with response). In Figure 6J, to determine whether percent RS change can differ depending on response types, a two sample *t* test was performed between neurons with onset responses only (n = 30 neurons) vs. neurons with onset followed by sustained responses (n = 35 neurons). In Figure 6K, whether the correlation coefficient between the best frequencies and the changes in RS was significantly different from zero was determined using a *t* test (n = 65). In Figure 6M, whether the population median frequency tuning width change was significantly different from zero was determined using a one sample Wilcoxon signed rank test (n = 60). In Figure 6L-N, neurons that lost all excitatory responses in the walking condition, yielding the tuning width of zero, were excluded (n = 5).

In Figure 7E, percent change in RS was compared across the three different spontaneous activity modulation categories using one-way ANOVA (of 65 neurons analyzed for evoked response, 39 showed increase, 8 showed decrease, and 18 showed no change in spontaneous activity). In Figure 7F, percent change in firing rates during response window were compared between RS (baseline subtracted) and <r> (without baseline subtraction) in each of the three spontaneous modulation categories. These comparisons were made using *t* tests only in neurons that showed significant attenuation in RS (red: n = 27 attenuated neurons of 39 neurons with increased spontaneous activity; blue: n = 6 attenuated neurons of 8 neurons with decreased spontaneous activity; gray: n = 14 attenuated neurons of 18 neurons with no change in spontaneous activity). Whether the evoked activity was significantly modulated by locomotion was determined using a two sample *t* test on RS values from stationary and walking trials. A neuron was considered modulated if there was a significant modulation in RS at least at one of the tone frequencies. In Figure 7G, to determine whether the correlation coefficient between the modulation of spontaneous rate and the evoked rate (<r>) was significantly different from zero, *t* tests were performed in the group with increased (n = 39) or decreased spontaneous activity (n = 8).

## Acknowledgements

This study was supported by the Institute for Basic Science in Korea (IBS-R015-D1). We thank Hyesook Lee for her help with histology, Dr. Seong-Gi Kim for helpful discussions and support, Drs. Karl Kandler and Mimi Kao for their critical comments on earlier versions of the manuscript.

## References

Adams JC. 1979. Ascending projections to the inferior colliculus. J Comp Neurol 183:519–538. doi: 10.1002/cne.901830305

Adams JC. 1980. Crossed and descending projections to the inferior colliculus. Neurosci Lett 19:1–5. doi:10.1016/0304-3940(80)90246-3

Aitkin LM, Dickhaus H, Schult W, Zimmermann M. 1978. External nucleus of inferior colliculus: auditory and spinal somatosensory afferents and their interactions. J Neurophysiol 41:837–847. doi:10.1152/jn.1978.41.4.837

Aitkin LM, Kenyon CE, Philpott P. 1981. The representation of the auditory and somatosensory systems in the external nucleus of the cat inferior colliculus. J Comp Neurol 196:25–40. doi:10.1002/cne.901960104

Aizenberg M, Mwilambwe-Tshilobo L, Briguglio JJ, Natan RG, Geffen MN. 2015. Bidirectional regulation of innate and learned behaviors that rely on frequency discrimination by cortical inhibitory neurons. PLOS Biology 13:e1002308. doi:10.1371/journal.pbio.1002308

Bigelow J, Morrill RJ, Dekloe J, Hasenstaub AR. 2019. Movement and VIP interneuron activation differentially modulate encoding in mouse auditory cortex. eNeuro ENEURO. 0164-19.2019. doi:10.1523/ENEURO.0164-19.2019

Bock GR, Webster WR. 1974. Coding of spatial location by single units in the inferior colliculus of the alert cat. Exp Brain Res 21:387–398. doi:10.1007/BF00237901

Caggiano V, Leiras R, Goñi-Erro H, Masini D, Bellardita C, Bouvier J, Caldeira V, Fisone G, Kiehn O. 2018. Midbrain circuits that set locomotor speed and gait selection. Nature 553:455–460. doi:10.1038/nature25448

Carcea I, Insanally MN, Froemke RC. 2017. Dynamics of auditory cortical activity during behavioural engagement and auditory perception. Nat Commun 8:14412. doi:10.1038/ncomms14412

Chen C, Cheng M, Ito T, Song S. 2018. Neuronal Organization in the Inferior Colliculus Revisited with Cell-Type-Dependent Monosynaptic Tracing. J Neurosci 38:3318–3332. doi:10.1523/JNEUROSCI.2173-17.2018

Clause A, Nguyen T, Kandler K. 2011. An acoustic startle-based method of assessing frequency discrimination in mice. Journal of Neuroscience Methods 200:63–67. doi: 10.1016/j.jneumeth.2011.05.027

Coleman JR, Clerici WJ. 1987. Sources of projections to subdivisions of the inferior colliculus in the rat. J Comp Neurol 262:215–226. doi:10.1002/cne.902620204

Cooper MH, Young PA. 1976. Cortical projections to the inferior colliculus of the cat. Experimental Neurology 51:488–502. doi:10.1016/0014-4886(76)90272-7

Creutzfeldt O, Ojemann G, Lettich E. 1989. Neuronal activity in the human lateral temporal lobe. Exp Brain Res 77:476–489. doi:10.1007/BF00249601

Dadarlat MC, Stryker MP. 2017. Locomotion Enhances Neural Encoding of Visual Stimuli in Mouse V1. J Neurosci 37:3764–3775. doi:10.1523/JNEUROSCI.2728-16.2017

Doupe AJ. 1997. Song-and Order-Selective Neurons in the Songbird Anterior Forebrain and their Emergence during Vocal Development. J Neurosci 17:1147–1167. doi:10.1523/JNEUROSCI.17-03-01147.1997

Egorova M, Ehret G, Vartanian I, Esser K-H. 2001. Frequency response areas of neurons in the mouse inferior colliculus. I. Threshold and tuning characteristics. Exp Brain Res 140:145–161. doi:10.1007/s002210100786

Ehret G. 1975. Frequency and intensity difference limens and nonlinearities in the ear of the housemouse (Mus musculus). J Comp Physiol 102:321–336. doi:10.1007/BF01464344

Eliades SJ, Wang X. 2013. Comparison of auditory-vocal interactions across multiple types of vocalizations in marmoset auditory cortex. J Neurophysiol 109:1638–1657. doi: 10.1152/jn.00698.2012

Escabí MA, Higgins NC, Galaburda AM, Rosen GD, Read HL. 2007. Early cortical damage in rat somatosensory cortex alters acoustic feature representation in primary auditory cortex. Neuroscience 150:970–983. doi: 10.1016/j.neuroscience.2007.07.054

Escabí MA, Schreiner CE. 2002. Nonlinear Spectrotemporal Sound Analysis by Neurons in the Auditory Midbrain. J Neurosci 22:4114–4131. doi:10.1523/JNEUROSCI.22-10-04114.2002

Farley GR, Morley BJ, Javel E, Gorga MP. 1983. Single-unit responses to cholinergic agents in the rat inferior colliculus. Hearing Research 11:73–91. doi: 10.1016/0378-5955(83)90046-1

Goyer D, Silveira MA, George AP, Beebe NL, Edelbrock RM, Malinski PT, Schofield BR, Roberts MT. 2019. A novel class of inferior colliculus principal neurons labeled in vasoactive intestinal peptide-Cre mice. eLife. doi:10.7554/eLife.43770

Groh JM, Trause AS, Underhill AM, Clark KR, Inati S. 2001. Eye Position Influences Auditory Responses in Primate Inferior Colliculus. Neuron 29:509–518. doi:10.1016/S0896-6273(01)00222-7

Gruters KG, Groh JM. 2012. Sounds and beyond: multisensory and other non-auditory signals in the inferior colliculus. Front Neural Circuits 6. doi:10.3389/fncir.2012.00096

Guo W, Clause AR, Barth-Maron A, Polley DB. 2017. A Corticothalamic Circuit for Dynamic Switching between Feature Detection and Discrimination. Neuron 95:180–194.e5. doi: 10.1016/j.neuron.2017.05.019

Hormigo S, e Horta Junior J de A de C, Gómez-Nieto R, López García DE. 2012. The selective neurotoxin DSP-4 impairs the noradrenergic projections from the locus coeruleus to the inferior colliculus in rats. Front Neural Circuits 6. doi:10.3389/fncir.2012.00041

Hoz L de, Nelken I. 2014. Frequency Tuning in the Behaving Mouse: Different Bandwidths for Discrimination and Generalization. PLOS ONE 9:e91676. doi:10.1371/journal.pone.0091676

Huffman RF, Henson OW. 1990. The descending auditory pathway and acousticomotor systems: connections with the inferior colliculus. Brain Res Brain Res Rev 15:295–323.

Ito S, Feldheim DA, Litke AM. 2017. Segregation of Visual Response Properties in the Mouse Superior Colliculus and Their Modulation during Locomotion. J Neurosci 37:8428–8443. doi:10.1523/JNEUROSCI.3689-16.2017

Ito T, Bishop DC, Oliver DL. 2009. Two Classes of GABAergic Neurons in the Inferior Colliculus. J Neurosci 29:13860–13869. doi:10.1523/JNEUROSCI.3454-09.2009

Jain R, Shore S. 2006. External inferior colliculus integrates trigeminal and acoustic information: Unit responses to trigeminal nucleus and acoustic stimulation in the guinea pig. Neuroscience Letters 395:71–75. doi:10.1016/j.neulet.2005.10.077

Jen PH-S, Sun X, Chen QC. 2001. An electrophysiological study of neural pathways for corticofugally inhibited neurons in the central nucleus of the inferior colliculus of the big brown bat, Eptesicus fuscus. Experimental Brain Research 137:292–302. doi:10.1007/s002210000637

Kanold PO, Young ED. 2001. Proprioceptive Information from the Pinna Provides Somatosensory Input to Cat Dorsal Cochlear Nucleus. J Neurosci 21:7848–7858. doi: 10.1523/JNEUROSCI.21-19-07848.2001

Klepper A, Herbert H. 1991. Distribution and origin of noradrenergic and serotonergic fibers in the cochlear nucleus and inferior colliculus of the rat. Brain Research 557:190–201. doi:10.1016/0006-8993(91)90134-H

Künzle H. 1998. Origin and terminal distribution of the trigeminal projections to the inferior and superior colliculi in the lesser hedgehog tenrec. European Journal of Neuroscience 10:368–376. doi:10.1046/j.1460-9568.1998.00020.x

Lee AM, Hoy JL, Bonci A, Wilbrecht L, Stryker MP, Niell CM. 2014. Identification of a Brainstem Circuit Regulating Visual Cortical State in Parallel with Locomotion. Neuron 83:455–466. doi:10.1016/j.neuron.2014.06.031

Lesica NA, Grothe B. 2008. Dynamic Spectrotemporal Feature Selectivity in the Auditory Midbrain. J Neurosci 28:5412–5421. doi:10.1523/JNEUROSCI.0073-08.2008

Lesica NA, Lingner A, Grothe B. 2010. Population Coding of Interaural Time Differences in Gerbils and Barn Owls. J Neurosci 30:11696–11702. doi:10.1523/JNEUROSCI.0846-10.2010

Lesicko AMH, Hristova TS, Maigler KC, Llano DA. 2016. Connectional Modularity of Top-Down and Bottom-Up Multimodal Inputs to the Lateral Cortex of the Mouse Inferior Colliculus. J Neurosci 36:11037–11050. doi:10.1523/JNEUROSCI.4134-15.2016

Li H, Mizuno N. 1997. Single neurons in the spinal trigeminal and dorsal column nuclei project to both the cochlear nucleus and the inferior colliculus by way of axon collaterals: a fluorescent retrograde double-labeling study in the rat. Neurosci Res 29:135–142. doi: 10.1016/s0168-0102(97)00082-5

Malmierca MS. 2004. The Inferior Colliculus: A Center for Convergence of Ascending and Descending Auditory Information. NBA 3:215–229. doi:10.1159/000096799

McGinley MJ, David SV, McCormick DA. 2015. Cortical Membrane Potential Signature of Optimal States for Sensory Signal Detection. Neuron 87:179–192. doi:10.1016/j.neuron.2015.05.038

Mineault PJ, Tring E, Trachtenberg JT, Ringach DL. 2016. Enhanced Spatial Resolution During Locomotion and Heightened Attention in Mouse Primary Visual Cortex. J Neurosci 36:6382–6392. doi:10.1523/JNEUROSCI.0430-16.2016

Morest DK, Oliver DL. 1984. The neuronal architecture of the inferior colliculus in the cat: defining the functional anatomy of the auditory midbrain. J Comp Neurol 222:209–236. doi:10.1002/cne.902220206

Motts SD, Schofield BR. 2009. Sources of cholinergic input to the inferior colliculus. Neuroscience 160:103–114. doi:10.1016/j.neuroscience.2009.02.036

Nelson A, Mooney R. 2016. The Basal Forebrain and Motor Cortex Provide Convergent yet Distinct Movement-Related Inputs to the Auditory Cortex. Neuron 90:635–648. doi: 10.1016/j.neuron.2016.03.031

Nelson A, Schneider DM, Takatoh J, Sakurai K, Wang F, Mooney R. 2013. A Circuit for Motor Cortical Modulation of Auditory Cortical Activity. J Neurosci 33:14342–14353. doi: 10.1523/JNEUROSCI.2275-13.2013

Niell CM, Stryker MP. 2010. Modulation of visual responses by behavioral state in mouse visual cortex. Neuron 65:472–479. doi:10.1016/j.neuron.2010.01.033

Oertel D, Young ED. 2004. What’s a cerebellar circuit doing in the auditory system? Trends in Neurosciences 27:104–110. doi:10.1016/j.tins.2003.12.001

Olthof BM, Rees A, Gartside SE. 2019. Multiple non-auditory cortical regions innervate the auditory midbrain. J Neurosci 1436–19. doi:10.1523/JNEUROSCI.1436-19.2019

Ono M, Bishop DC, Oliver DL. 2017. Identified GABAergic and Glutamatergic Neurons in the Mouse Inferior Colliculus Share Similar Response Properties. J Neurosci 37:8952–8964. doi:10.1523/JNEUROSCI.0745-17.2017

Ono M, Oliver DL. 2014. The Balance of Excitatory and Inhibitory Synaptic Inputs for Coding Sound Location. J Neurosci 34:3779–3792. doi:10.1523/JNEUROSCI.2954-13.2014

Porter KK, Metzger RR, Groh JM. 2007. Visual-and saccade-related signals in the primate inferior colliculus. PNAS 104:17855–17860. doi:10.1073/pnas.0706249104

Porter KK, Metzger RR, Groh JM. 2006. Representation of Eye Position in Primate Inferior Colliculus. Journal of Neurophysiology 95:1826–1842. doi:10.1152/jn.00857.2005

Poulet JFA, Hedwig B. 2002. A corollary discharge maintains auditory sensitivity during sound production. Nature 418:872. doi:10.1038/nature00919

Reimer J, McGinley MJ, Liu Y, Rodenkirch C, Wang Q, McCormick DA, Tolias AS. 2016. Pupil fluctuations track rapid changes in adrenergic and cholinergic activity in cortex. Nature Communications 7:13289. doi:10.1038/ncomms13289

Robards MJ. 1979. Somatic neurons in the brainstem and neocortex projecting to the external nucleus of the inferior colliculus: an anatomical study in the opossum. J Comp Neurol 184:547–565. doi:10.1002/cne.901840308

Rockel AJ, Jones EG. 1973. The neuronal organization of the inferior colliculus of the adult cat. II. The pericentral nucleus. J Comp Neurol 149:301–334. doi:10.1002/cne.901490303

Rummell BP, Klee JL, Sigurdsson T. 2016. Attenuation of Responses to Self-Generated Sounds in Auditory Cortical Neurons. J Neurosci 36:12010–12026. doi:10.1523/JNEUROSCI.1564-16.2016

Schneider DM, Mooney R. 2018. How Movement Modulates Hearing. Annual Review of Neuroscience 41:553–572. doi:10.1146/annurev-neuro-072116-031215

Schneider DM, Nelson A, Mooney R. 2014. A synaptic and circuit basis for corollary discharge in the auditory cortex. Nature 513:189–194. doi:10.1038/nature13724

Schneider DM, Sundararajan J, Mooney R. 2018. A cortical filter that learns to suppress the acoustic consequences of movement. Nature 561:391–395. doi:10.1038/s41586-018-0520-5

Schnupp JW, King AJ. 1997. Coding for auditory space in the nucleus of the brachium of the inferior colliculus in the ferret. J Neurophysiol 78:2717–2731. doi:10.1152/jn.1997.78.5.2717

Schuller G. 1979. Vocalization influences auditory processing in collicular neurons of the CF-FM-bat, Rhinolophus ferrumequinum. J Comp Physiol 132:39–46. doi:10.1007/BF00617730

Shore SE. 2005. Multisensory integration in the dorsal cochlear nucleus: unit responses to acoustic and trigeminal ganglion stimulation. European Journal of Neuroscience 21:3334–3348. doi:10.1111/j.1460-9568.2005.04142.x

Singla S, Dempsey C, Warren R, Enikolopov AG, Sawtell NB. 2017. A cerebellum-like circuit in the auditory system cancels responses to self-generated sounds. Nat Neurosci 20:943–950. doi:10.1038/nn.4567

Suga N, Schlegel P. 1972. Neural attenuation of responses to emitted sounds in echolocating rats. Science 177:82–84. doi:10.1126/science.177.4043.82

Tammer R, Ehrenreich L, Jürgens U. 2004. Telemetrically recorded neuronal activity in the inferior colliculus and bordering tegmentum during vocal communication in squirrel monkeys (Saimiri sciureus). Behavioural Brain Research 151:331–336. doi:10.1016/j.bbr.2003.09.008

Winer JA, Schreiner CE. 2005. The Inferior Colliculus. New York, Springer.

Williamson RS, Hancock KE, Shinn-Cunningham BG, Polley DB. 2015. Locomotion and Task Demands Differentially Modulate Thalamic Audiovisual Processing during Active Search. Current Biology 25:1885–1891. doi:10.1016/j.cub.2015.05.045

Woolley SMN, Portfors CV. 2013. Conserved mechanisms of vocalization coding in mammalian and songbird auditory midbrain. Hear Res 305:45–56. doi:10.1016/j.heares.2013.05.005

Xiong XR, Liang F, Li H, Mesik L, Zhang KK, Polley DB, Tao HW, Xiao Z, Zhang LI. 2013. Interaural Level Difference-Dependent Gain Control and Synaptic Scaling Underlying Binaural Computation. Neuron 79:738–753. doi:10.1016/j.neuron.2013.06.012

Xiong XR, Liang F, Zingg B, Ji X, Ibrahim LA, Tao HW, Zhang LI. 2015. Auditory cortex controls sound-driven innate defense behaviour through corticofugal projections to inferior colliculus. Nat Commun 6:7224. doi:10.1038/ncomms8224

Zhou J, Shore S. 2006. Convergence of spinal trigeminal and cochlear nucleus projections in the inferior colliculus of the guinea pig. Journal of Comparative Neurology 495:100–112. doi:10.1002/cne.20863

Zhou M, Liang F, Xiong XR, Li L, Li H, Xiao Z, Tao HW, Zhang LI. 2014. Scaling down of balanced excitation and inhibition by active behavioral states in auditory cortex. Nature Neuroscience 17:841–850. doi:10.1038/nn.3701

